# Dissociated face- and word-selective intracerebral responses in the human ventral occipito-temporal cortex

**DOI:** 10.1101/2020.12.23.423819

**Authors:** Simen Hagen, Aliette Lochy, Corentin Jacques, Louis Maillard, Sophie Colnat-Coulbois, Jacques Jonas, Bruno Rossion

## Abstract

The extent to which faces and written words share neural circuitry in the human brain is actively debated. Here we compared face-selective and word-selective responses in a large group of patients (N = 37) implanted with intracerebral depth electrodes in the ventral occipito-temporal cortex (VOTC). Both face-selective (i.e., significantly different responses to faces vs. nonface visual objects) and word-selective (i.e., significantly different responses to words vs. pseudofonts) neural activity is isolated through frequency-tagging. Critically, this sensitive approach allows to objectively quantify category-selective neural responses and disentangle them from general visual responses. About 70% of significant contacts show either only face-selectivity or only word-selectivity, with the expected right and left hemispheric dominance, respectively. Spatial dissociations are also found within core regions of face and word processing, with a medio-lateral dissociation in the fusiform gyrus (FG) and surrounding sulci, while a postero-anterior dissociation is found in the inferior occipital gyrus (IOG). Only 30% of the significant contacts show both face- and word-selective responses. Critically, in these contacts, across the VOTC or in the FG and surrounding sulci, between-category selective-amplitudes (faces vs. words) showed no-to-weak correlations, despite strong correlations in both the within-category selective amplitudes (face-face, word-word) and the general visual responses to words and faces. Overall, we conclude that category-selectivity for faces and written words is largely dissociated in the human VOTC.

**Significance Statement:** In modern human societies, faces and written words have become arguably the most significant stimuli of the visual environment. Despite extensive research in neuropsychology, electroencephalography and neuroimaging over the past three decades, whether these two types of visual signals are recognized by similar or dissociated processes and neural networks remains unclear. Here we provide an original contribution to this outstanding scientific issue by directly comparing frequency-tagged face- and word-selective neural responses in a large group of epileptic patients implanted with intracerebral electrodes covering the ventral occipito-temporal cortex. While general visual responses to words and faces show significant overlap, the respective category-selective responses are neatly dissociated in spatial location and magnitude, pointing to largely dissociated processes and neural networks.

## Introduction

The human ventral occipito-temporal cortex (VOTC) is crucial for visual recognition. Bilateral or right unilateral VOTC damage can cause a detrimental recognition impairment specific to faces (prosopagnosia, Bodamer, 1947; Meadows, 1974), while a selective lesion to the left occipitotemporal region can produce specific written word recognition impairment (pure alexia; Cohen et al., 2004; see Farah, 1991). These findings are complemented by functional magnetic resonance magnetic (fMRI) studies, showing that different VOTC regions respond selectively to faces and words. Most notably, a category-selective response to faces is found in the lateral parts of the middle fusiform gyrus and in the inferior occipital gyrus, with a right hemispheric dominance (Puce et al., 1995; Kanwisher et al., 1997; Haxby et al., 2000; Grill-Spector et al., 2017 for review) while written words evoke a category-selective response in the left posterior fusiform and occipitotemporal sulcus (Grill-Spector & Weiner, 2014; Devlin et al., 2006). While these findings support the claim that the VOTC contains dissociated neural circuitry for face and written word recognition, fMRI studies showing partial spatial overlap between the functional face- and word-selective regions, in particular the fusiform gyrus (e.g., Dehaene et al., 2010; Harris et al., 2016), have probed researchers to speculate instead that faces and words share the same neural circuitry and functional processes (Behrmann and Plaut, 2020; Behrmann and Plaut, 2013; Nestor et al., 2012; Robinson et al., 2017).

Understanding the neural processes that generate recognition of visual words and faces is of high relevance for human social communication, because for most humans in literate societies, faces and written words constitute, arguably, the two most common, complex and socially relevant categories of their everyday visual environment. The claim that these fundamental behaviors are supported by same neural processes is controversial and has implications for the large portion of the visual neuroscience community who studies either face or word recognition. Importantly, this issue extends beyond the realm of faces and words, and speaks directly to how the brain organize to perform stimulus-response mappings to highly experienced and behaviorally relevant visual categories in the two hemispheres, an issue cutting straight to the core of human neuroscience.

To shed original light on this issue, we provide the first comprehensive and systematic comparison of face and word category-selective neural responses, using intracerebral (i.e., cortical depth electrodes) recordings across the VOTC of a large group of epileptic patients. These recordings provide direct measures of local cortical activity, allowing to explore up to the anterior sections of the VOTC without being affected by magnetic susceptibility artifacts as in fMRI (Rossion et al., 2018; Wandell, 2011). The patients were tested here with two frequency-tagging experiments to isolate face- and word-selective responses. In the face condition, variable object images appear at a fixed frequency rate (F) with variable face-images interleaved as every 5^th^ item (F/5), while in the word condition, variable pseudofonts appear at a fixed frequency rate (F) with variable words interleaved as every 5th image (Figure 1). As shown in scalp EEG and intracerebral recording studies, this approach objectively quantifies intracerebral face- and word-selective responses at the face- and word-stimulation F/5 frequencies and harmonics (Rossion et al., 2015; Jonas et al., 2016; Lochy et al., 2018). Given that the same electrode contacts are recorded in the same individual brains in the two paradigms, including an extensive sampling of anterior VOTC regions, this approach is particularly well suited to test the issue of spatial and functional overlap of face- and word-selective neural activity in the VOTC. Importantly, the frequency tagging paradigm allows for isolating category-*selective* responses at the F/5 frequency and parse out *general* visual responses at the base rate F frequencies (Rossion et al., 2018). Thus, if face and words share neuro-functional circuitry, one should observe a large overlap of *selective* responses obtained after parsing out shared general visual responses, as well as a high correlation of response amplitude across significant contacts for the two categories.

**Figure 1.**
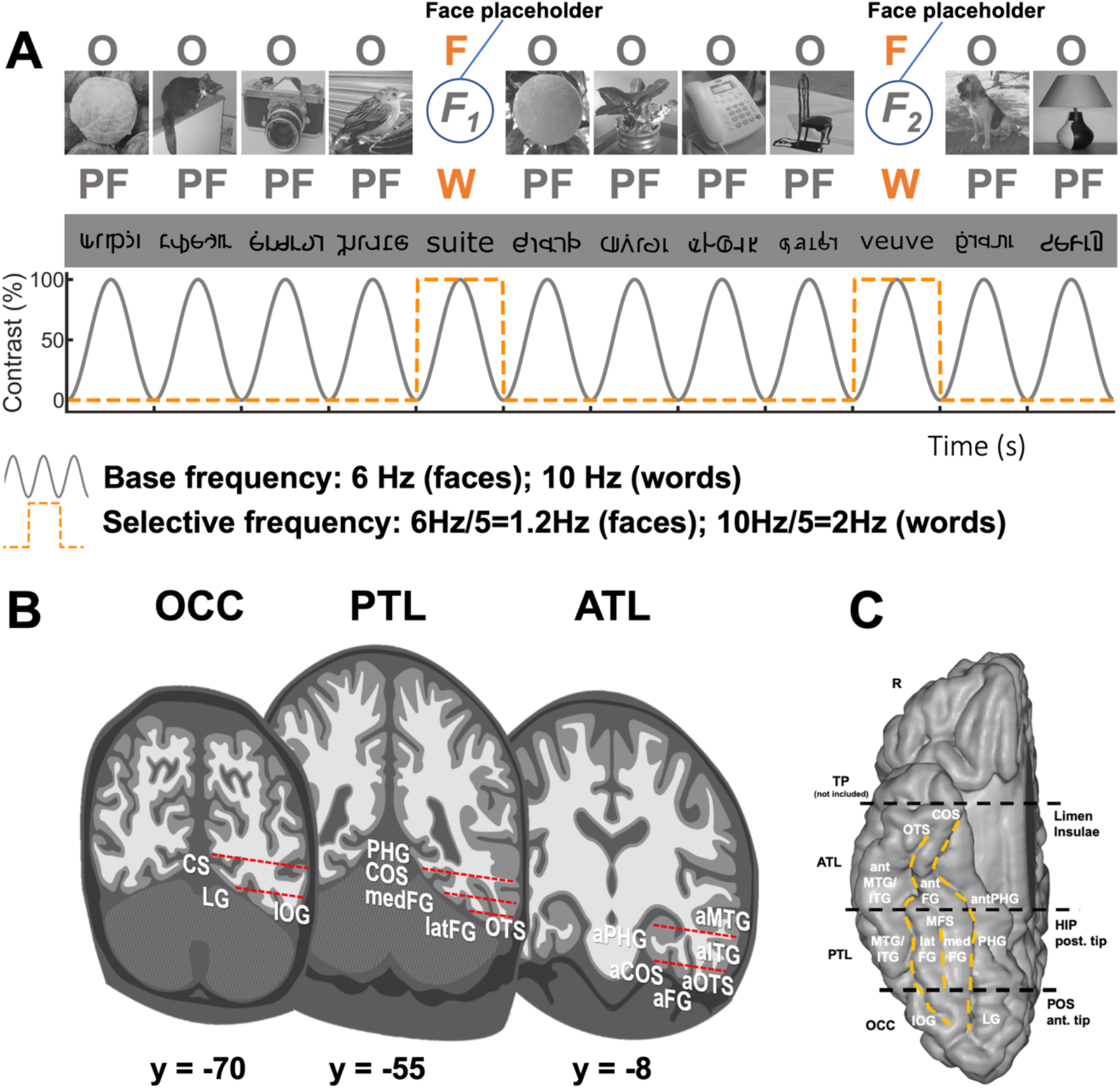
FPVS and SEEG methods. **A.** For faces, images of living or non-living objects were presented by sinusoidal contrast modulation at a rate of 6 stimuli per second (6 Hz) with different images of faces presented in separate sequences every 5 stimuli (i.e., appearing at the frequency of 6 Hz/5 = 1.2 Hz) (Faces are here indicated by placeholder F; see Jonas et al., 2016 for examples). For words, pseudofonts (PF) were presented at a rate of 10 stimuli per second (10 Hz) with different words presented in separate sequences every 5 stimuli (i.e., appearing at the frequency of 10 Hz/5 = 2 Hz). **B.** Schematic coronal representation of the typical trajectories of depth electrodes implanted in the VOTC (adapted from Jonas et al., 2016; Lochy et al., 2018). Electrodes consist of 8-15 contiguous recording contacts (red rectangles) spread along the electrode length, along the medio-lateral axis. **C.** Schematic representation of the parcellation scheme used to determine the anatomical label of each contact. Anatomical regions were defined in each individual hemisphere according to major anatomical landmarks. The ventral temporal sulci (COS, OTS, and midfusiform sulcus, i.e., MFS) serve as medial/lateral borders of regions, whereas two coronal reference planes containing anatomical landmarks (posterior tip of the hippocampus, i.e., HIP and anterior tip of the parieto-occipital sulcus, i.e., POS) serve as an anterior/posterior boundary for each region. We considered contacts in the ATL if they were located anteriorly to the posterior tip of the hippocampus. Note that we did not include in our analyses contacts in the temporal pole (TP), i.e., anterior to the limen insulae. The schematic locations of these anatomical structures are shown on a reconstructed cortical surface of the Colin27 brain. Acronyms: ATL: anterior temporal lobe; PTL: posterior temporal lobe; OCC: occipital lobe; PHG: parahippocampal gyrus; COS: collateral sulcus; FG: fusiform gyrus; ITG: inferior temporal gyrus; MTG: middle temporal gyrus; OTS: occipito-temporal sulcus; CS: calcarine sulcus; IOG: inferior occipital gyrus; LG: lingual gyrus; a: anterior; lat: lateral; med: medial.

## Methods

### Participants

The study included 37 patients (21 females, mean age: 32.9 ± 8.4 years, 36 right-handed) undergoing clinical intracerebral evaluation with depth electrodes (StereoElectroEncephaloGraphy, SEEG) for refractory partial epilepsy, studied in the Epilepsy Unit of the University Hospital of Nancy between September 2013 and June 2016. Patients were included in the study if they had at least one intracerebral electrode implanted in the VOTC (Figure 1B). The SEEG data for word stimulation from 36 patients were included in Lochy et al. (2018), while the data for face stimulation from all patients were included in Hagen et al. (2020). All patients gave written consent to participate to the study, which was part of a protocol approved by the Ethics committee of the University Hospital of Nancy.

### Intracerebral electrode implantation and recording

Intracerebral electrodes were stereotactically implanted within the participants’ brains for clinical purposes, i.e., to delineate their seizure onset zones and to functionally map the surrounding cortex in the perspective of an eventual epilepsy surgery (Bédos-Ulvin et al., 2017). Each 0.8 mm diameter intracerebral electrode contains 8-15 independent recording contacts of 2 mm in length separated by 1.5 mm from edge to edge (for details about the electrode implantation procedure, see Salado et al., 2017). Intracerebral EEG was sampled at a 512 Hz with a 256-channel amplifier and referenced to either a midline prefrontal scalp electrode (FPz, in 32 participants) or, when scalp electrodes were not available, an intracerebral contact in the white matter (6 participants). Similar to previous intracranial reports, the data in subsequent analysis were not re-referenced (e.g., Allison, McCarthy, 2002; Hagen et al., 2020; Jonas et al. 2016; Kadipasaogly et al. 2016; Lochy et al., 2018; Matsuo et al., 2015; Nobre, McCarthy, 1995). However, a separate analysis using a bipolar re-reference yielded the same data pattern. Noise from ocular and muscle activity is of little worry with the frequency tagging approach since the broadband noise is smeared across a large number of frequency noise bins (faces = 32256; words = 32768), while the signal is contained in a few narrow bins (faces = 12; words = 4) (Regan, 1989). Contacts located in brain lesions visible on structural MRI were excluded from any analysis. The recorded sequences were checked by an expert epileptologist (author JJ), and sequences with epileptic discharges or epileptic seizures were removed from the analysis. Note that sequences with epileptic spikes were kept because epileptic spikes do not occur at the exact same frequency of occurrence that the frequency of interest, and is therefore spread across a large number of noise bins. To assess the effect of epileptic spikes on data, a previous report with the same approach applied artefact rejection (removing epileptic spikes), whereby it was shown that the pattern of results did not change (Jonas et al., 2016, appendix).

### Fast periodic visual stimulation paradigm

#### Stimuli

In the face condition, we used 200 grayscale natural images of various non-face objects (from 14 non-face categories: cats [n=9], dogs [n=5], horses [n=5], birds [n=24], flowers [n=15], fruits [n=28], vegetables [n=21], houseplants [n=15], phones [n=13], chairs [n=15], cameras [n=6], dishes [n=15], guitars [n=15], lamps [n=14]) and 50 grayscale natural images of faces (all images taken from a paradigm validated in scalp EEG studies, e.g., Rossion et al., 2015). Each image contained an unsegmented object or a face near the center. Faces and objects varied substantially in terms of size, viewpoint, lighting conditions and background across images (see Rossion et al., 2015). Images were equalized for mean pixel luminance and contrast (i.e. standard deviation across pixels).

In the word condition, we used words and pseudofonts (30 of each type), all composed of five elements (letters or pseudofonts, PF). These stimuli were also taken from a paradigm validated in scalp EEG studies, e.g., Lochy et al., 2015). French words were selected from the Lexique 3.55 database (New et al., 2001) with the following criteria: they were frequent common nouns (84.99 per million) in singular form, with limited orthographic neighbors (average, 1.9; range from 0 to 4), no foreign language origin, and no accents. PF items were built on an item-by-item basis: letters from words were vertically flipped, segmented, and segments were rearranged into five pseudoletters with the same overall size as the original word. Each word thus had a corresponding PF containing the exact same amount of black-on-white contrast, so that all conditions were similar in terms of lower-level visual properties. Bigram frequencies were calculated with Wordgen (Duyck et al., 2004) and are reported as summated type bigram frequencies (from the French CELEX database). Stimuli were presented in Verdana font, with the size ranging from 4.8 to 7.7 (width) and 1.15 to 2 (height) degrees of visual angle.

#### Experimental procedure

In the face condition, participants viewed continuous sequences of natural images of objects presented at a fast rate of 6 Hz through sinusoidal contrast modulation, in which faces were presented periodically as every 5th stimulus so that the frequency of face presentation was 1.2 Hz (i.e. 6 Hz/5) (Figure 1, Movie S1). All images were randomly selected from their respective categories. For the word condition, participants viewed continuous sequences of pseudofont strings (PF) presented periodically at a rate of 10 Hz through sinusoidal contrast modulation (from 0 to 100% in 50ms, then back to 0% in 50 ms) with randomly selected words inserted every fifth item, so that the word presentation frequency was 2 Hz (10 Hz/5) (Figure 1A and Movies S2). In all conditions, a sequence lasted 70 s: 66 s of stimulation at full-contrast flanked by 2 s of fade-in and fade-out, where contrast gradually increased or decreased, respectively. During the sequences, participants performed a color-change detection task on the fixation cross. In the face condition, they were instructed to detect brief (500 ms) black to red changes, while in the word condition, the fixation cross was blue and changed to red. In the face condition, sequences were repeated a minimum of two times (average sequences across patients: 2.67, range: 2-6). In the word condition, the experiment was repeated a minimum of two times for all but two patients (average sequences across patients: 2.92, range: 1-6). *Control procedure.* In a control condition with houses, participants viewed continuous sequences of natural images of objects presented periodically at 6 Hz through sinusoidal contrast modulation, with randomly selected house images inserted every fifth item, so that the frequency of house presentation was 1.2 Hz (i.e. 6 Hz/5) (see Figure 1 in Hagen et al., 2020). Thus, the procedure was identical to faces with the exception that variable natural house images was interspersed at the fifth cycle rather than images of faces. All images were randomly selected from their respective categories. Note that a full report of face-house intracerebral responses, as measured with FPVS-SEEG, have been reported elsewhere (Hagen et al., 2020), and that here we use the overlap between faces and houses strictly as a control comparison to face-word-overlap.

Participants were not informed about the periodicity of the stimulation and were unaware of the objectives of the study. No participant had seizures in the 2 hours preceding fast periodic visual stimulation (FPVS) recordings.

#### Frequency domain processing

Signals corresponding to the face and words conditions were processed the same way. Segments of SEEG corresponding to stimulation sequences were extracted (74-second segments, −2 s to +72 s). The 74 s data segments were cropped to contain an integer number of 1.2 Hz cycles (for faces) and 2 Hz cycles (for words) beginning 2 s after the onset of the sequence (right at the end of the fade-in period) until approximately 65 s (for faces) and 66 s (for words), i.e. before stimulus fade-out (75 face cycles ≈ 63 s; 126 word cycles ≈ 63 s). No artifact correction was applied because (1) (S)EEG artifacts generate noise at frequencies that locate mostly outside of the frequencies of interest (1.2 Hz or 2.0 Hz and associated harmonics) and, most importantly, the noise is broadband while the signal locates in narrow frequency bins due to the very high frequency resolution of our approach (Regan, 1989; Rossion, 2014). Hence, analyzing such data with or without artifact rejection leads to the same results (Jonas et al., 2016). Sequences of recorded voltage (i.e., time-domain) were averaged separately for each participant and condition. Averaging sequences in the time-domain before the Fast Fourier Transform (FFT) increases signal-to-noise ratio by cancelling out non-phase locked noise. Subsequently, an FFT was applied to the full length of the cropped averaged time sequences. The amplitude spectra were extracted for all contacts by taking the modulus of the Fourier coefficients at each frequency bin normalized (by dividing) by half of the number of time samples in the time series. The long recording sequence resulted in a spectrum with a high frequency resolution of 0.0159 (1/63sec) for both faces and words (thus, faces and words did not differ in terms of frequency resolution). No data segments were excluded from the analysis. No other processing was performed to the data.

#### Selective responses

The FPVS approach used here allows identifying and separating two distinct types of responses in both conditions: (1) a general visual response occurring at the base stimulation frequency (faces: 6 Hz; words: 10 Hz) and its harmonics, as well as (2) a category-selective response at 1.2 Hz (faces) and 2 Hz (words) and its harmonics (face-selective or word-selective response, also called face categorization or word categorization response, respectively). In both face and word conditions, category-selective responses significantly above noise level at the face/word frequency (1.2 Hz / 2 Hz) and its harmonics were determined as follows: (1) the FFT spectrum was cut into 4 segments centered at the face/word frequency and harmonics, from the 1st until the 4th (faces: 1.2 Hz until 4.8 Hz; words: 2 Hz until 8 Hz), and surrounded by 25 neighboring bins on each side (Figure 2A); (2) the amplitude values in these 4 segments of FFT spectra were summed (Figure 2B); (3) the summed FFT spectrum was transformed into a Z-score (Figure 2C). Z-scores were computed as the difference between the amplitude at the face/word frequency bin and the mean amplitude of 48 surrounding bins (25 bins on each side, excluding the 2 bins directly adjacent to the bin of interest, i.e., 48 bins) divided by the standard deviation of amplitudes in the corresponding 48 surrounding bins. A contact was considered as showing a selective response in a given condition if the Z-score at the frequency bin of face or word stimulation exceeded 3.1 (i.e., *p* < 0.001 one-tailed: signal > noise).

**Figure 2.**
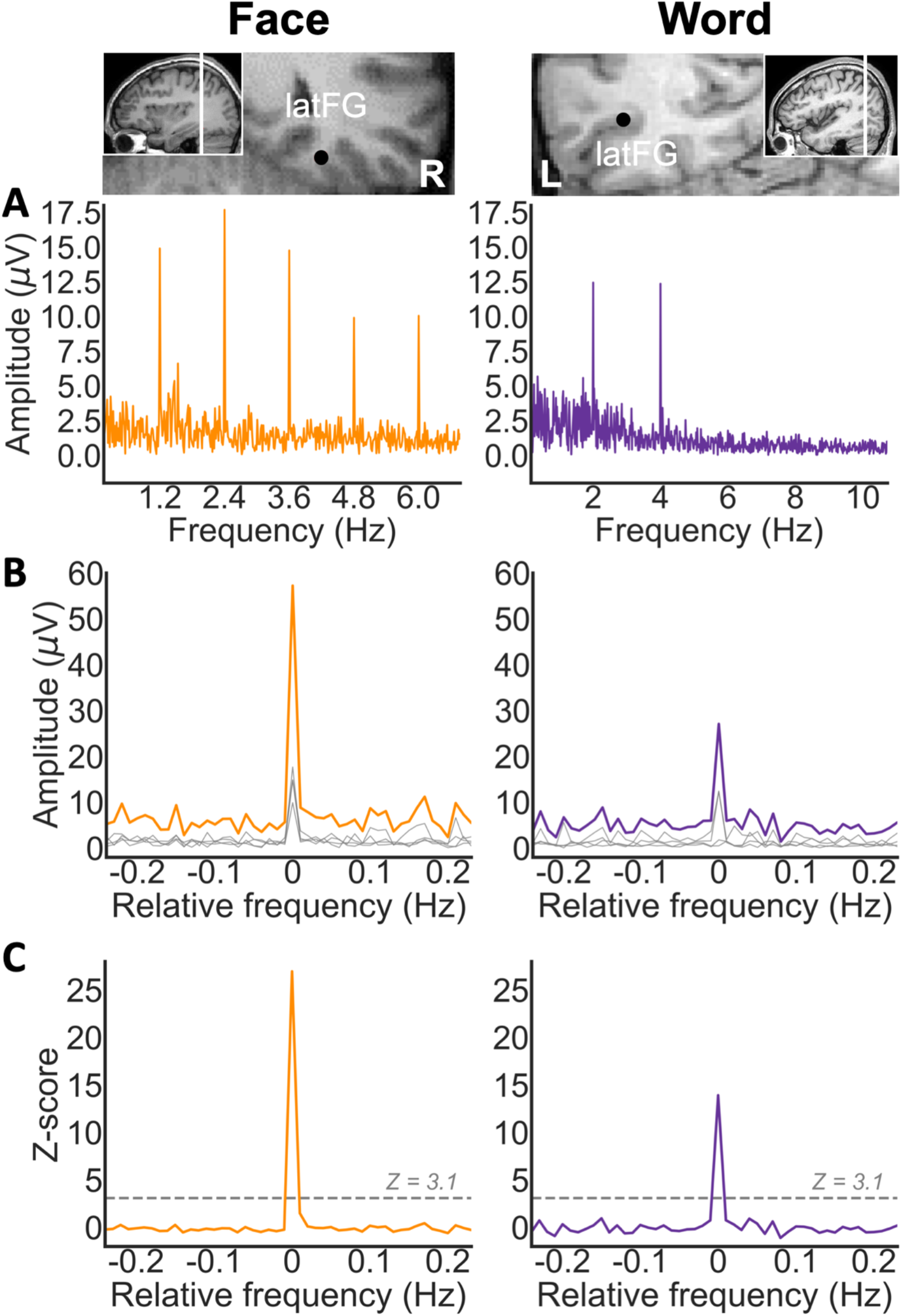
Intracerebral selective responses recorded in the VOTC. **A.** Intracerebral EEG frequency-domain responses recorded at an individual recording contact (raw FFT amplitude) located in the right latFG of a single participant during a face stimulation sequence, and in the left latFG in a single participant during a word stimulation sequence. The anatomical location of the contact is shown in a coronal MRI slice. Face-selective responses are observed at 1.2 Hz and harmonics and word-selective responses are observed at 2 Hz and harmonics. **B**. Significant face- and word-selective responses were determined by first segmenting the FFT spectrum into four segments centered at the frequency of face- and word-stimulation and its harmonics up to the 4th harmonic (i.e., faces: 1.2, 2.4, 3.6, and 4.8 Hz; words: 2, 4, 6, and 8 Hz). Individual FFT segments are shown in gray (see horizontal gray bars on the X axis in A, representing the length of each FFT segment). The four segments, containing both the signal and the surrounding noise, were then summed (orange and purple lines for faces and words, respectively). The 0 mark corresponds to the face or word stimulation frequencies. **C.** Z-score transformation of the summed FFT spectrum for statistical purpose. The Z score at the face/word frequency exceeds 3.1 (p < 0.001), indicating that these contact show significant face- or word-selective responses.

#### Classification of significant contacts

Based on the pattern of discrimination responses across the 2 conditions (i.e., significant or not), we labeled each significant contact as follows: (i) contacts showing a significant face categorization response, but not a significant word categorization response, were defined as “face” (+face, g-word); (ii) contacts showing a significant word response, but not a significant face categorization response, were defined as “word” (−face, +word); and (iii) contacts showing significant categorization responses to both faces and words, were defined as “overlap” (+face, +word).

#### Quantification of response amplitude

Baseline-corrected amplitudes were computed as the difference between the amplitude at each frequency bin and the average of 48 corresponding surrounding bins (up to 25 bins on each side, i.e., 50 bins, excluding the 2 bins directly adjacent to the bin of interest, i.e., 48 bins). Face-selective responses were quantified separately as the sum of the baseline-subtracted amplitudes at the face frequency from the 1st until the 14th (1.2 Hz until 16.8 Hz), excluding the 5th and 10th harmonics (6 Hz and 12 Hz) that coincided with the base rate frequency (Jonas et al., 2016). Word-selective responses were quantified separately as the sum of the baseline-subtracted amplitudes at the face frequency from the 1st until the 4th (2 Hz until 8 Hz; Lochy et al., 2018). The range of harmonics used was related to the highest harmonic with a significant response (Jonas et al., 2016; Lochy et al., 2018). Base rate response amplitudes were quantified separately as the sum of the baseline-subtracted amplitudes at the base frequency from the 1st until the 4th (faces: 6 Hz until 24 Hz; words: 10 Hz until 40 Hz), separately for face and words sequences. It is important to note that different frequencies were used for faces (1.2 Hz; 6 Hz) and words (2 Hz; 10 Hz) because they were deemed optimal for evoking *selective* responses in prior investigations examining the domains in isolation, thus maximizing the amount of overlap possibly detected (Jonas et al., 2016; Lochy et al., 2018).

### Contact Localization in the Individual Anatomy

The exact position of each contact in the individual anatomy was determined by fusing the postoperative CT scan with a T1-weighted MRI. Contacts inside the gray matter were anatomically labeled in the individual anatomy using the same topographic VOTC parcellation as in Lochy et al. (2018; Figure 1C) based on anatomical landmarks. Major VOTC sulci (collateral sulcus and occipito-temporal sulcus) served as medio-lateral divisions. Postero-anterior divisions were the anterior tip of the parieto-occipital sulcus for the border between occipital and temporal lobes, and the posterior temporal lobe (PTL) and the anterior temporal lobe (ATL). In addition, we created an anatomical region-of-interest (ROI) consisting of the fusiform gyrus and surrounding sulci that are considered core to face and word processing (e.g., Kanwisher et al., 1997; Cohen et al., 2001; Harris et al., 2016; for review, see Grill-Spector and Weiner, 2014). We refer to this region as FG+sulci, which according to our parcellation scheme (Figure S1), included latFG+OTS, medFG+CoS, posterior antOTS, antFG, and posterior antCOS (Y Talairach < −25).

### Proportion and amplitude maps in Talairach space

In a separate analysis, anatomical MRIs were spatially normalized to determine Talairach coordinates of intracerebral contacts. Electrode locations were transformed to Talairach space by first locating the contacts in the original MRI system. Next, using Advanced Source Analysis (ASA, https://www.nitrc.org/projects/asa/), we determined three anatomical landmarks (nasion, left and right pre-auricular points) in order to define the fiducial system. Then, several points where determined in the MR volume (e.g., anterior and posterior commissure), to introduce the Talairach system, which was a piecewise linear transformation of the AC-PC system: anterior and posterior point (AP and PP; i.e. point of the cortex with maximum and minimum x coordinates), superior and inferior points (SP and IP; i.e. point of the cortex with maximum and minimum z coordinates), and right and left points (RP and LP; i.e. point of the cortex with maximum and minimum y coordinates) (Koessler et al., 2009, appendix). Talairach coordinates of the intracerebral contacts were used to perform group analyses and visualization. The cortical surface used to display group maps was obtained from segmentation of the Colin27 brain from AFNI (Cox, 1996) which is aligned to the Talairach space. Using Talairach coordinates, we computed the local proportion and amplitudes of the discrimination intracerebral contacts across the VOTC. Local proportion and amplitudes of contacts were computed in volumes (i.e., “voxels”) of size 15 × 15 × 200 mm (for the X, left–right; Y, posterior–anterior; and Z, inferior–superior dimensions, respectively) by steps of 3 × 3 × 200 mm over the whole VOTC. A large voxel size in the Z dimension was used to collapse across contacts along the inferior–superior dimension. For each voxel, we extracted the following information across all participants in our sample: (i) the number of recorded contacts located within the voxel; (ii) the number of contacts showing a significant response for each type of discrimination; and (iii) the mean amplitudes in the significant contacts. For each voxel and each type of discrimination (i.e. face, word, overlap), we computed the proportion of significant contacts over recorded contacts (proportions are crucial here since sampling differs across regions), as well as the mean amplitudes over/in the significant contacts. To ensure reliability and reproducibility, we only considered voxels in which at least two participants showed significant responses. Then, for each voxel, we determined whether the proportion/amplitudes of significant contacts was significantly above zero using a bootstrap procedure in the following way: (i) sampling contacts from the voxel (the same number as the number of recorded contacts in the voxel) with replacement; (ii) determining the proportion of significant contacts for this bootstrap sample and storing this value; (iii) repeating steps i and ii 5,000 times to generate a distribution of bootstrap proportions and to estimate the *p*-value as the fraction of bootstrap proportions equal to zero.

### Correlation analysis

To compute within-category (e.g., face vs. face) and between-category (e.g., face vs. word) correlations, for each overlap-contact, two within-condition (e.g., face sequence 1 and 3) or between-condition sequences (face sequence 1 and word sequence 2) were randomly sampled, respectively. To account for different combinations of selections, this procedure was repeated 5000 times for each hemisphere, thereby creating 5000 sample correlations with associated values for *t*, *p*, and 95% confidence intervals (*CI*s). Finally, a grand average across the 5000 values (*r, t, p, 95%CIs*) was computed. Computing correlations based on sampling two sequences per intracerebral recording contact ensures that both the within- and between-category correlations are based on an equal amount of data (e.g., an alternative approach of splitting the within-category data would result in within-category correlations that was computed with half the amount of data compared to between-category correlations).

A permutation approach was used to statistically test correlations. First, to test for overall statistical significance (R > 0), (1) each of the 5000 sample correlations was recomputed and stored, after randomly shuffling the order of one data vector (to disrupt structure in the data), (2) this procedure was repeated 1000 times to produce a sampling distribution reflecting the null hypothesis, (3) estimate a P-value as the fraction of permuted correlations larger than or equal to the real/measured correlation (two-tailed). Second, to test for statistical difference relative to another correlation (e.g., R1 [face-face] vs. R2 [face-word] > 0), (1) each of the 5000 sample correlations for the two correlations (R1 and R2) was recomputed after combining, shuffling, and splitting half the data from each correlation (e.g., R1 [Face1-Face2] vs. R2 [Face3-Word1]: combine, shuffle, and split Face2 and Word1), (2) subtract and store the recomputed sample correlations to produce 5000 permuted difference correlations, (3) repeat this procedure 1000 times to produce a sampling null distribution, (4) estimate a P-value as the fraction of permuted correlations larger than or equal to the difference of the real/measured correlations (one-tailed).

## Results

### Visual face and word categorization

We found 566 contacts with category-selective responses (for faces, and/or words) in 36 individual brains, that is 28.75% of total recorded contacts (1969 contacts implanted in the grey matter of the VOTC in 37 subjects). Among these contacts, 15.90% were selective to words *only* (“word” contacts, 90/566, subjects = 22), 53.36% were selective to faces *only* (“face” contacts, 302/566, subjects = 33). Interestingly, 30.74% contacts were selective to both words and faces (“face-word-overlap” contacts, 174/566, subjects = 24). An example of the response profile of each contact type is shown in Figure 3. Each contact was localized in the individual anatomy using a topographic parcellation of the VOTC and in the Talairach space to perform group analyses and visualization (Figure 3; see methods and Figure 1C for the description of the parcellation; Table 1 for contact count by hemisphere and region).

**Figure 3.**
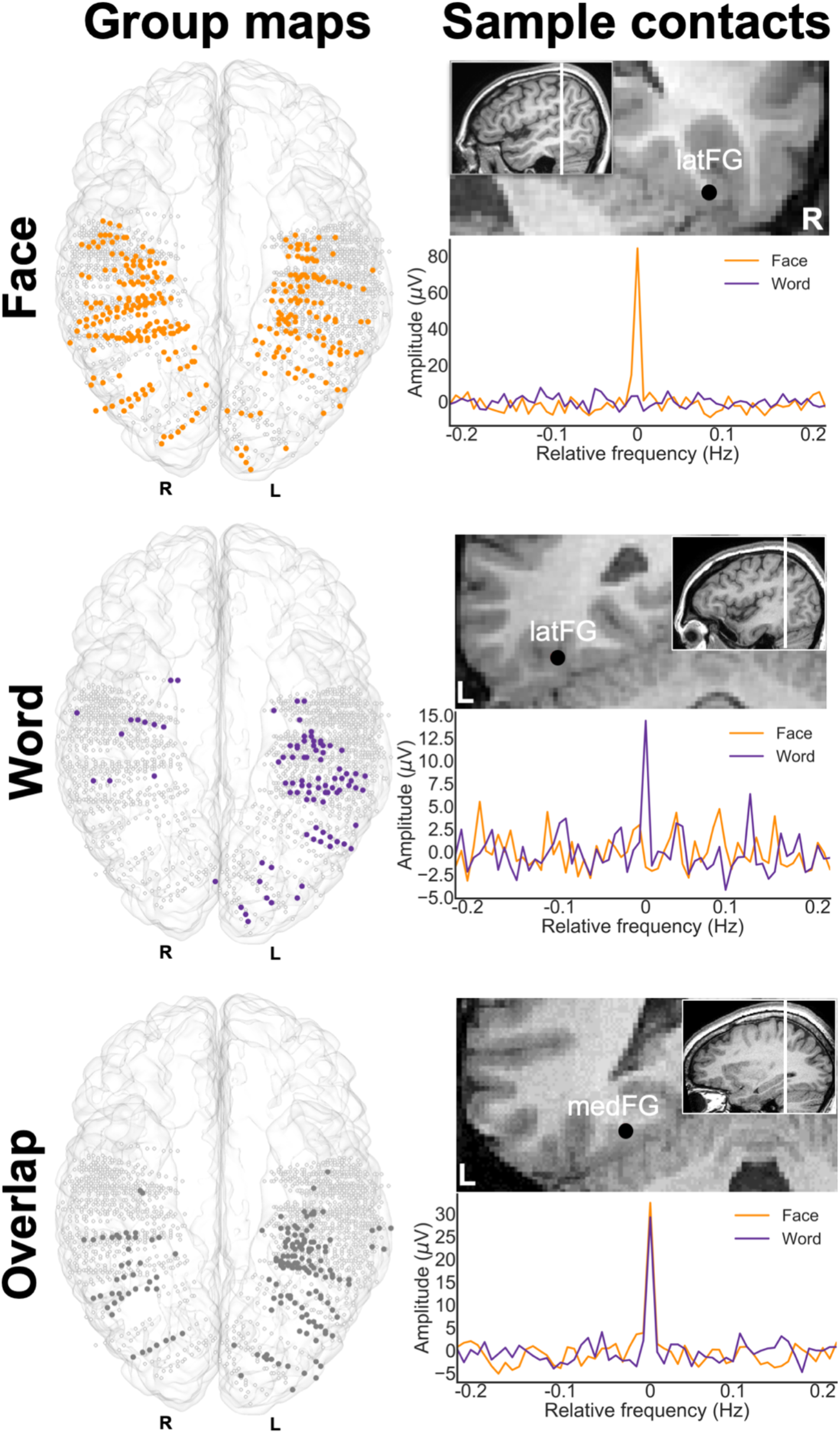
Classification and distribution of three types of category-selective contacts. **Left.** Maps of all 1969 VOTC recording contacts across the 37 individual brains displayed in the TAL space using a transparent reconstructed cortical surface of the Colin27 brain. Each circle represents a single contact. Color filled circles correspond to significant contacts colored according to their category selectivity (face, word, face-word-overlap). White-filled circles correspond to contacts on which no selective responses were recorded. **Right.** Examples of baseline corrected FFT spectra for each contact type (3 individual contacts in 3 different participants). Top: face contact, selective to faces only, middle: word contact selective to words only; bottom: face-word-overlap contact selective to both faces and words. Their anatomical location is illustrated in the respective coronal MRI slices.

**Table 1.** Face vs Words. Number of selective contacts and corresponding number of participants (in parentheses) in each anatomical region. Acronyms: ATL: anterior temporal lobe; PTL: posterior temporal lobe; OCC: occipital lobe; VMO: ventromedial occipital; IOG: inferior occipital gyrus; PHG: parahippocampal gyrus; FG: fusiform gyrus; MTG: middle temporal gyrus; ITG: inferior temporal gyrus; OTS: occipito-temporal sulcus; COS: collateral sulcus; ant: anterior; lat: lateral; med: medial.

|  | Left Hemisphere |  |  | Right Hemisphere |  |  |
| --- | --- | --- | --- | --- | --- | --- |
| Regions | Faces | Words | Overlap | Faces | Words | Overlap |
| VMO | 14 (5) | 11 (5) | 17 (5) | 20 (4) | 0 (0) | 4 (1) |
| IOG | 6 (4) | 8 (2) | 19 (5) | 12 (3) | 0 (0) | 9 (3) |
| **Total OCC** | **20 (9)** | **19 (7)** | **36 (10)** | **32 (7)** | **0 (0)** | **13 (4)** |
| PHG | 2 (1) | 0 (0) | 0 (0) | 1 (1) | 0 (0) | 0 (0) |
| medFG | 24 (10) | 4 (3) | 27 (12) | 19 (6) | 1 (1) | 4 (2) |
| latFG | 8 (5) | 12 (6) | 21 (8) | 24 (6) | 1 (1) | 7 (3) |
| MTG/ITG | 14 (6) | 10 (6) | 5 (3) | 10 (4) | 1 (1) | 4 (1) |
| **TOTAL PTL** | **48 (22)** | **26 (15)** | **53 (23)** | **54 (17)** | **3 (3)** | **15 (6)** |
| antPHG | 0 (0) | 0 (0) | 1 (1) | 0 (0) | 0 (0) | 0 (0) |
| antCOS | 29 (13) | 17 (10) | 8 (5) | 25 (10) | 6 (3) | 3 (3) |
| antFG | 2 (1) | 0 (0) | 10 (5) | 13 (5) | 0 (0) | 3 (2) |
| antOTS | 20 (11) | 9 (7) | 16 (9) | 22 (9) | 2 (2) | 5 (2) |
| antMTG/ITG | 9 (4) | 7 (2) | 7 (2) | 28 (10) | 1 (1) | 4 (2) |
| **Total ATL** | **60 (29)** | **33 (19)** | **42 (22)** | **88 (34)** | **9 (6)** | **15 (9)** |

### Spatially and functionally dissociated face-and word-selective responses

To visualize and quantify face and word contacts at a group level, local proportions (out of total recorded contacts) and local average amplitudes (in significant contacts) were computed and projected on the cortical surface (Figure 4). Contacts that were selective only to faces (face contacts) or only to words (word contacts) accounted for most of the total significant contacts (392/566 = 69.26%). Out of these contacts, there were more face than word contacts (*diff* = (302/566) (F) – (90/566) (W) = 37.46%; *χ*^*2*^(1, *N* = 566) = 175.39, *p* < 0.001). Moreover, the mean face-selective amplitude in face contacts (*M* = 19.02 μV) was larger than the mean word-selective amplitude in word contacts (*M* = 8.66 μV, *Mdiff* = 10.36 μV, *t(390)* = 5.69, *p* < 0.001).

**Figure 4.**
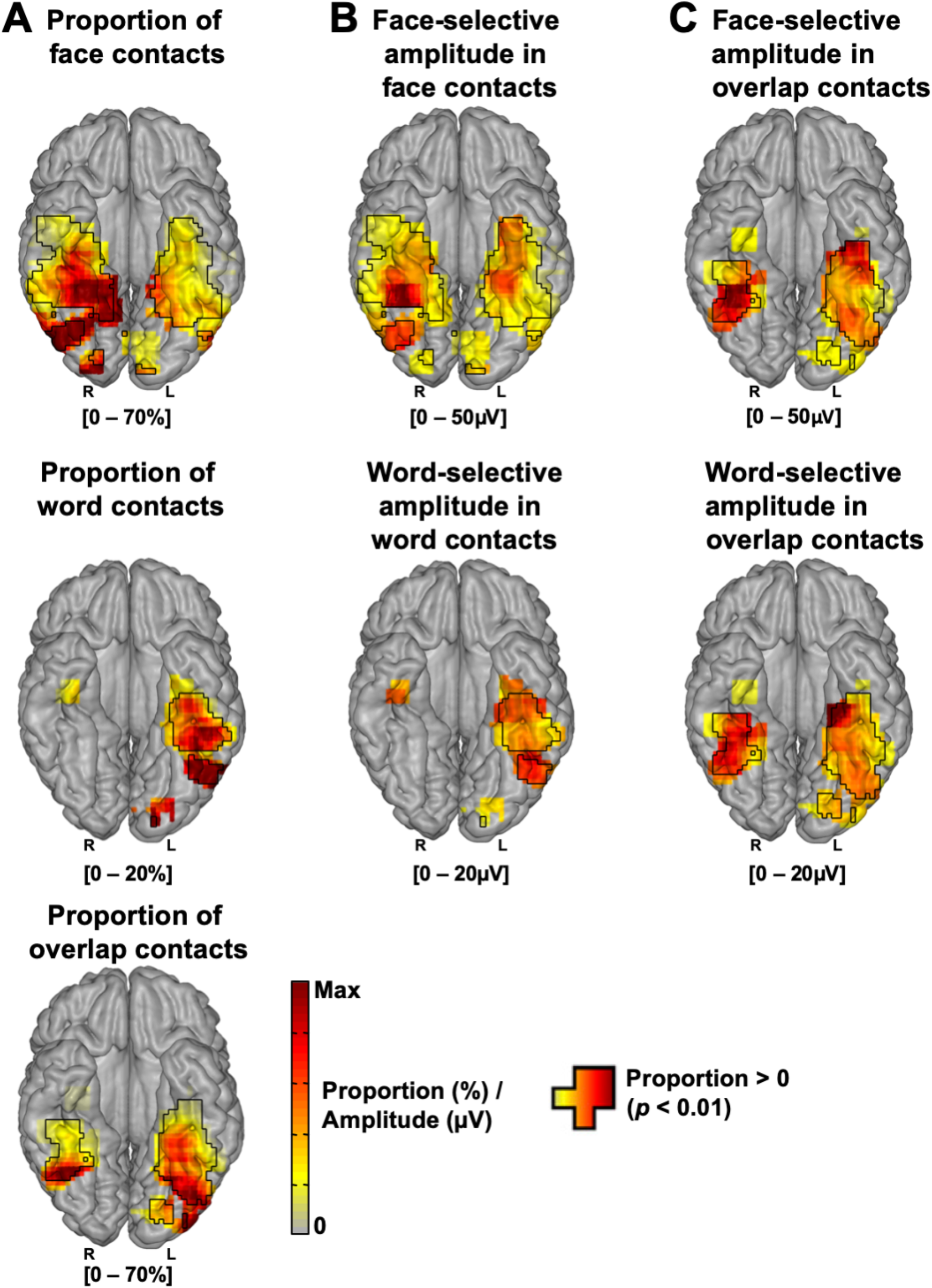
Proportion and amplitude maps. **A.** Maps of the local proportion of selective contacts relative to recorded contacts across VOTC for each contact type (face, word, overlap), displayed on the cortical surface. **B.** Maps of the local mean selective response amplitudes for face- and word-contacts. **C.** Maps of the local mean face- and word-selective response amplitudes in overlap contacts. Local proportions and amplitudes were computed in 15 × 15 voxels (for X and Y dimensions, respectively) using contacts collapsed over the Z dimension (superior–inferior) for better visualization. For the sake of replicability, only voxels containing significant responses from at least two individual brains were considered. Black contours outline proportions and amplitudes significantly above zero.

The face and word contacts showed the expected right and left hemispheric dominance, respectively. Specifically, faces made up a larger proportion of the significant contacts in the right than the left hemisphere (*diffhemi* = Right (R) – Left (L) = (174/229) - (128/337) = 38.00%, *χ*^*2*^(1, *N* = 566) = 79.11, *p* < 0.001; Figure 5A), pposite was true for words (*diffhemi* = (12/229) (R) - (78/337) (L) = −17.91%, *χ*^*2*^(1, *N* = 566) = 32.69, *p* < 0.001; Figure 5A). Moreover, for faces there was a trend towards higher face-selective amplitudes in the right (*M* = 20.61 μV) than the left hemisphere (*M* = 16.87 μV; *Mdiff* = 3.74 μV, *t(300)* = 1.89, *p* = 0.060; Figure 5B), while the low number of word contacts in the right hemisphere (Figure 5A; *n* right hemisphere = 12) prevented the equivalent comparison for words (see Figure 5B for point estimates).

**Figure 5.**
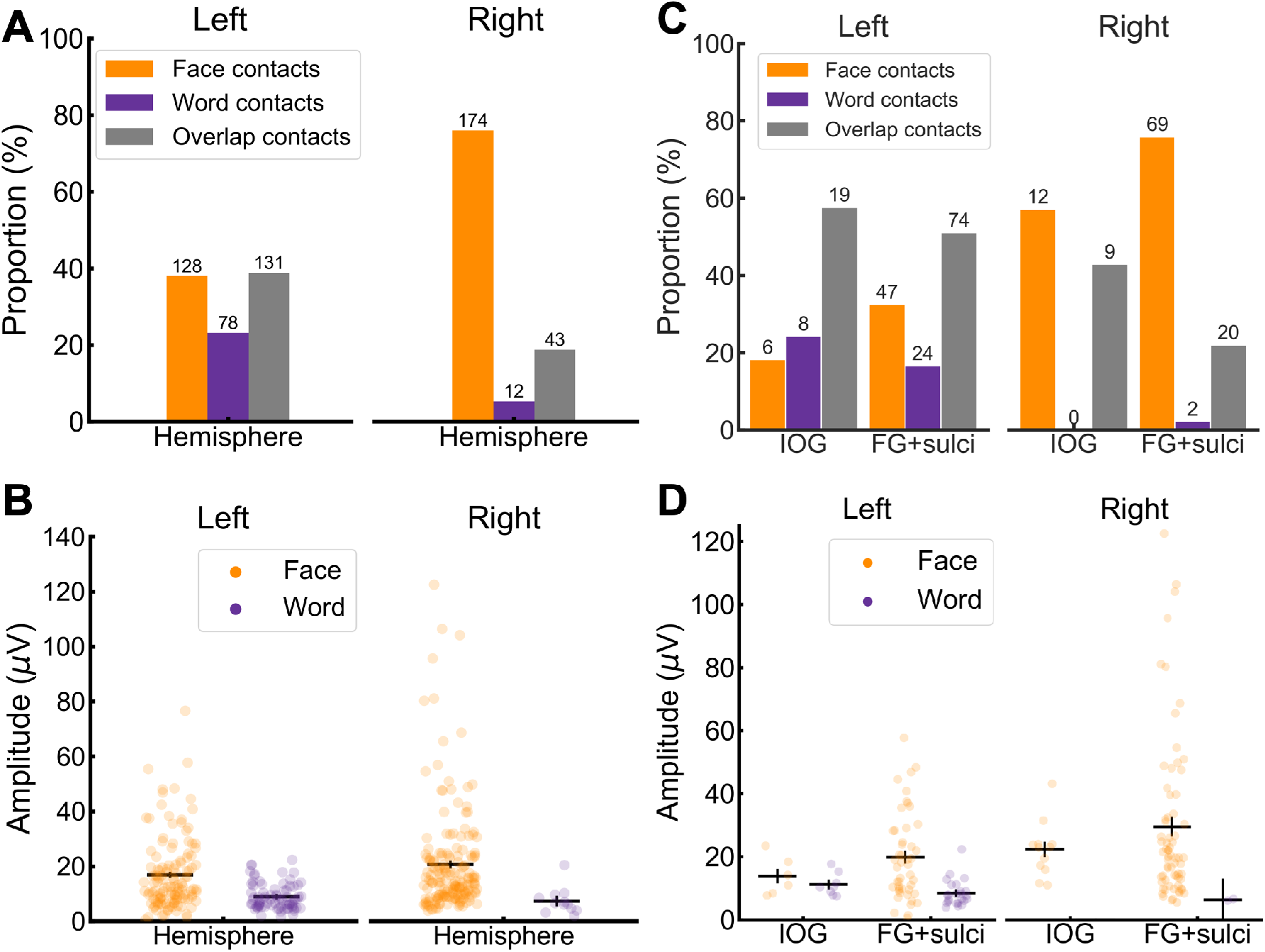
**A.** Proportion of significant contacts (out of total significant contacts) split by contact-type in each hemisphere. **B.** Average selective amplitude in significant contacts split by contact-type in each hemisphere. Each dot represents a single contact. **C.** Proportion of significant contacts (out of total significant contacts) split by contact-type in the IOG and FG+sulci. **D.** Average selective amplitude in significant contacts split by contact-type in the IOG and the FG+sulci. Same convention as panel B. The numbers on top of the bars in panel A and C indicate the number of significant contacts. Error bars in panels B and D represent standard error of the means.

The hemispheric dominance was also observed within anatomical regions thought to be core to face and word processing, the FG+sulci and IOG (e.g., Grill-Spector and Weiner, 2014; Davies-Thompson et al., 2016), and where it has been proposed that faces and words compete for and share neural processes (Nestor et al., 2012; Behrmann and Plaut, 2013). Specifically, in both the FG+sulci and the IOG, faces made up a larger proportion of significant contacts in the right than the left hemisphere (FG+sulci: *diffhemi* = (69/91) (R) - (47/145) (L) = 43.41%, *χ*^*2*^(1, *N* = 236) = 40.44, *p* < 0.001; IOG: *diffhemi* = (12/21) (R) - (6/33) (L) = 38.96%, *χ*^*2*^(1, *N* = 54) = 7.10, *p* = 0.008; Figure 5C), whereas the opposite was true for words (FG+sulci: *diffhemi* = (2/91) (R) - (24/145) (L) = −14.35%, *χ*^*2*^(1, *N* = 236) = 10.33, *p* = 0.001; IOG: *diffhemi* = (0/21) (R) - (8/33) (L) = −24.24%, *χ*^*2*^(1, *N* = 54) = 4.21, *p* = 0.040; Figure 5C). Similarly, in the FG+sulci and IOG, the face-selective amplitudes were larger in the right than the left hemisphere (*FG+sulci: Mdiff* = 29.50 μV (R) - 19.78 (L) = 9.72 μV, *t*(114) = 2.34, *p* = 0.021; IOG: *Mdiff* = 22.27 (R) - 13.82 (L) = 8.45 μV, *t*(16) = 2.10, *p* = 0.052; Figure 5D; note low number of IOG contacts), while the low number of word contacts in the right hemisphere (IOG = 0; FG+sulci = 2) prevented the equivalent comparison for words (Figure 5D).

Next, we analyzed the differences between face and word contacts in their central mass along the X and Y Talairach axis, which was restricted to the left hemisphere due to the small number of significant word contacts in the right hemisphere (*n* = 12; see Figure 4A second row; Table 1). Overall, the central mass of X Talairach coordinates (mean Talairach X) for word contacts was more lateral (*Mtalx* = 37.03 mm) than for the face contacts (*Mtalx* = 33.03 mm; *Mdiff* = 4 mm, *t*(df=204) = 2.25, *p* = 0.026; Figure 6A), while along the Y Talairach axis there was no difference between faces (*Mtaly* = −38.91 mm) and words (*Mtaly* = −43.51 mm; *Mdiff* = 4.60 mm, *t*(df=204) = 1.42, *p* = 0.159; Figure 6A). In the FG+sulci, there was also a medio-lateral dissociation (29.15 (F) – 35.46 (W) = −6.31 mm; *t*(df=69) = −3.53, *p* < 0.001; Figure 6B), but no posterior-anterior dissociation (−40.38 (F) – (−38.79) (W) = −1.59 mm; *t*(df=69) = −0.71, *p* = 0.480; Figure 6C). In contrast, in the IOG, there was no medio-lateral dissociation (45.00 (F) – 44. 63 (W) = 0.38 mm; *t*(df=12) = 0.12, *p* = 0.909; Figure 6B), but a posterior-anterior dissociation (−70.33 (F) – (−64.00) (W) = −6.33 mm; *t*(df=12) = −2.80, *p* = 0.016; note few contacts in IOG; Figure 6C). Thus, in addition to a large-scale dissociation along the medio-lateral axis, we found spatial dissociations within local regions that are core to face and word recognition.

**Figure 6.**
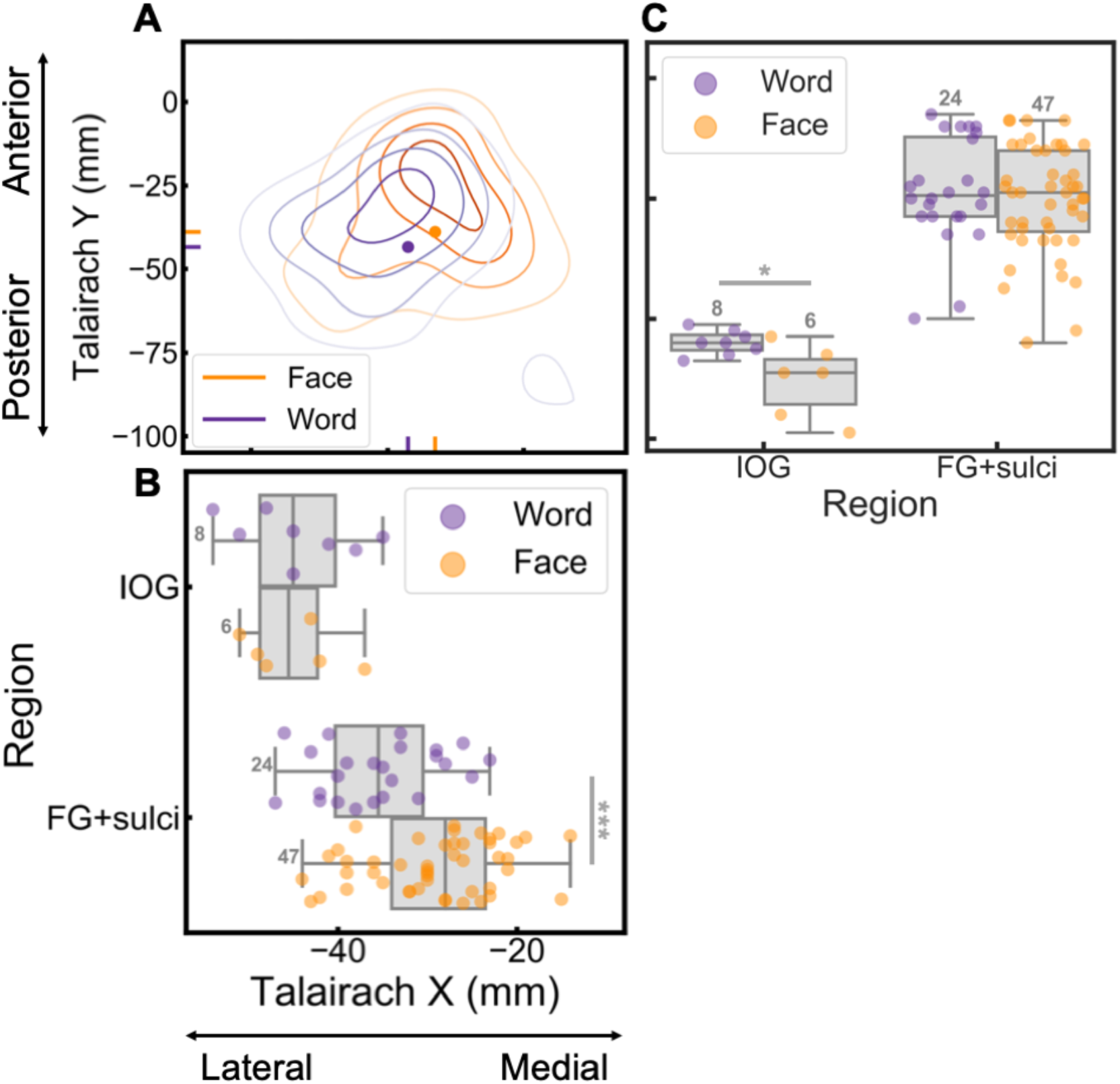
Spatial distribution of word and face contacts along medio-lateral (Talairach X) and posterio-anterior (Talairach Y) axis. The right hemisphere was not analyzed to due to it containing a low number of word contacts. **A.** Contour plot showing distribution of word and face contacts along the medio-lateral (Talairach X) and posterio-anterior axes (Talairach Y). Each distribution is computed relative to itself and darker contour colors indicate larger density of contacts. The central mass for each distribution is plotted as a solid dot within the contour plot and as a vertical line on the x- and y-axis. **B.** X coordinates of word and face contacts in core regions for neural processes supporting face and word recognition (IOG, FG+sulci). The boxplot displays three quartiles (Q1, median, Q2) and the whiskers extend to points that lie within 1.5 interquartile range (IQRs) of the lower and upper quartile. * p < 0.05; *** p < 0.001. The number next to the error bar indicate the number of contacts. Note that overlapping contacts are indicated by darker colors. **C.** Y coordinates of face and word contacts in the IOG and FG+sulci. Same convention as for panel B.

In summary, more than two-thirds of all significant intracerebral contacts were functionally dissociated, either responding selectively only to faces or only to words. Moreover, these functionally dissociated contacts showed spatial dissociations, including opposite hemispheric dominance and dissociations along the medio-lateral and posterior-anterior axis.

### Overlapping but functionally dissociated face- and word-selective responses

Notably, a considerable proportion of contacts were also responding selectively to *both* faces and words (Figure 3; Figure 5A; Table 1). Does this overlap truly reflect shared neural populations recruited during both face and word processing? Proponents of this view have argued that faces and words share neural populations due to shared representational demands (e.g., central-view representations; Behrmann and Plaut, 2013).

To test this view, we first we examined the quantity of face-word-overlap contacts in each hemisphere, the FG+sulci, and in the IOG. The face-word-overlap contacts accounted for a total of 30.74% (174/566) of all significant contacts and there was a larger proportion of overlap contacts (out of total significant contacts) in the left than in the right hemisphere (*diffhemi* = (43/229) (R) - (131/337) (L) = −20.10%, *χ*^*2*^ (1, *N* = 566) *=* 25.86, *p* < 0.001, Figure 7A). Most of the face-word-overlap contacts (122/174 = 70.12%) were found within anatomical regions thought to be core to face and word processes, the FG+sulci and the IOG (FG+sulci: 94/174 = 54.02%; IOG: 28/174 = 16.09%; FG+sulci+IOG = 70.11% of all face-word-overlap contacts). Within the FG+sulci there was a larger proportion (out of significant contacts) of face-word-overlap in the left than the right hemisphere (*diffhemi* = (20/91) (R) - (74/145) (L) = −29.06%, *χ*^*2*^(1, *N* = 236) = 18.50, *p <* 0.001; Figure 7B) while there was an equal proportion in left and right IOG (*diffhemi* = (9/21) (R) - (19/33) (L) = −14.72%, *χ*^*2*^(1, *N* = 54) = −0.60, *p =* 0.438; Figure 7B).

**Figure 7.**
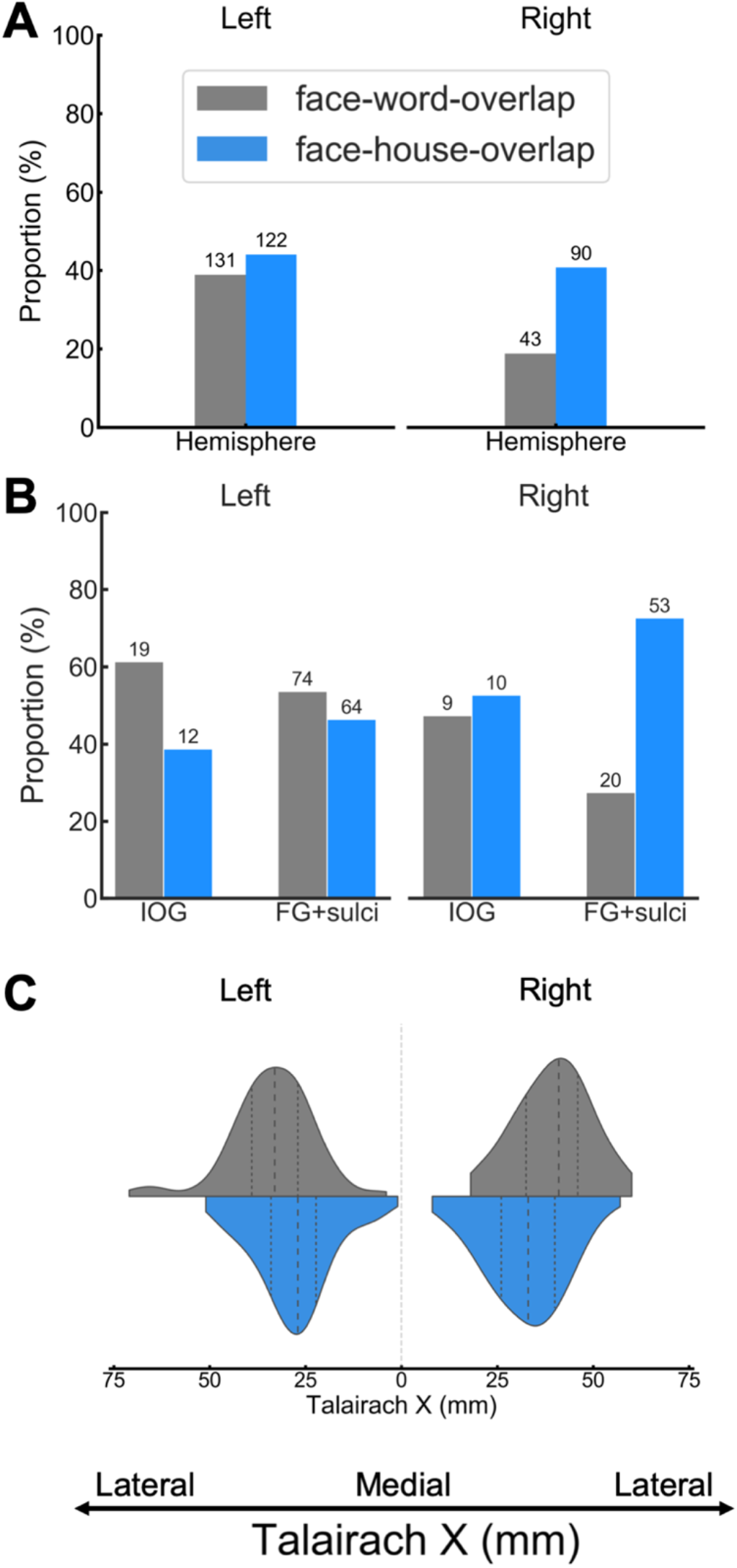
**A.** Proportion of face-word-overlap (of total significant contacts [face, word, overlap]) and proportion of face-house-overlap contacts (of total significant contacts [face, house, overlap]), split by hemisphere. The proportion is expressed in percentage (%). The number of significant contacts is indicated on the top of each bar. **B.** Proportion of overlap contacts, split by overlap-type, hemisphere and ROI. Same conventions as panel A. **C.** Distribution of overlap contacts, split by overlap-type and hemisphere, along the Talairach X axis. Dashed lines within each distribution reflects the median and upper and lower quartiles. The range of each distribution is cut to fit the range of the observed data (i.e., the gaussian density estimation does not assign probability to contact coordinates lower or higher than observed data).

To evaluate whether the overlap between face-selective and word-selective responses was specific to these two categories, houses were used as a control category. Since being landmark stimuli, pictures of houses are associated with dissociated representations from faces and words (e.g., peripheral-view rather than central-view representations; Plaut and Behrmann, 2011), and show a medio-lateral spatial dissociation with faces (e.g., Grill-Spector and Weiner, 2014). The house-selective responses were isolated similar to faces, by replacing faces with variable natural house images at the oddball rate of 1.2 Hz (see Methods for description of paradigm). Across all recorded contacts, contacts showing significant selective responses to both faces and houses were classified as face-house-overlap contacts (irrespective of how they responded to words). Note that a full description of face-house intracerebral responses, as measured with FPVS-SEEG, has been reported elsewhere (Hagen et al., 2020), and that here we use the overlap between faces and houses strictly as a comparison to face-word-overlap.

Notably, a substantial number of contacts showed both face-selective and house-selective responses (212/498 = 42.57% of significant face and house contacts). There was an equal proportion (out of significant face, house, overlap contacts) of overlap in across hemispheres (*diffhemi* = (90/221) (R) - (122/277) (L) = −3.32%, *χ*^*2*^(1, *N* = 448) *=* 0.43, *p* = 0.514, Figure 7A). Crucially, a substantial portion of these were also located within the IOG and the FG+sulci (FG+sulci: 117/212 = 55.19%; IOG: 22/212 = 10.38%; FG+sulci+IOG = 65.57% of all face-house-overlap contacts). There was an equal proportion of face-house-overlap contacts in the right and the left hemispheres of both the FG+sulci and the IOG (FG+sulci: *diffhemi* = (53/90) (R) - (64/127) (L) = 8.50%, *χ*^*2*^(1, *N* = 217) = 1.21, *p =* 0.272; IOG: *diffhemi* = (10/21) (R) - (12/29) (L) = 6.24%, *χ*^*2*^(1, *N* = 50) = 0.02, *p =* 0.881; Figure 7B). Thus, the spatial overlap between selective responses to faces and selective responses to words was not outstanding, since, if anything, a larger amount of overlap was also observed between faces and houses, both within hemispheres as well as within core face- and word-regions.

Next, we analyzed the face-word-overlap and face-house-overlap contacts along the X Talairach axis. If the overlap is a general phenomenon related to the resolution of neural recording, overlap-contacts should be situated close to the sources (in order for both to be captured in the same intracerebral recording contact). Thus, since words-selective responses are localized more laterally than face-selective responses, and house-selective responses are more medial than face-selective responses (e.g., fMRI: Grill-Spector and Weiner, 2014; Hasson et al., 2002; Spiridon et al., 2006; Nasr et al., 2011; intracranial EEG: Hagen et al., 2020; Kadipasaoglu et al., 2016; Jacques et al., 2016), the centre of mass of face-word-overlap contacts should be more lateral than the centre of mass of face-house-overlap contacts. Consistent with this claim, in both hemispheres, the centre of mass of the face-word-overlap contacts (Right: *Mtalx* = 39.54 mm; Left: *Mtalx* = 33.83 mm) was more lateral than that of the face-house-overlap contacts (Left: *Mtalxdiff* = 5.96 mm, *t*(251) = 4.51, *p* < 0.001; Right: *Mtalxdiff* = 7.11 mm, *t*(131) = 3.78, *p* < 0.001; Figure 7 C).

To directly test the extent of dissociation in the selective responses in the overlap contacts, we correlated face- and word-selective response amplitudes in face-word-overlap contacts, as well as the face- and house-selective responses in the face-house-overlap contacts. A shared neural population account would predict strong correlations between faces and words (which should be larger than that of faces and houses) since they are reflecting the same neural population. This was tested by correlating between-category response amplitudes across overlap contacts (i.e., faces vs. words; faces vs. houses), and comparing them to within-category correlations (i.e., faces vs. faces, words vs. words, houses vs. houses) in the same contacts. This was done separately by hemispheres (Figure 8A) and FG+sulci (Figure 8B), except in the IOG due to its low number of significant overlap contacts (face-words-overlap: *n* right = 9; *n* left = 19; face-houses-overlap: *n* right = 12; *n* left = 10).

**Figure 8.**
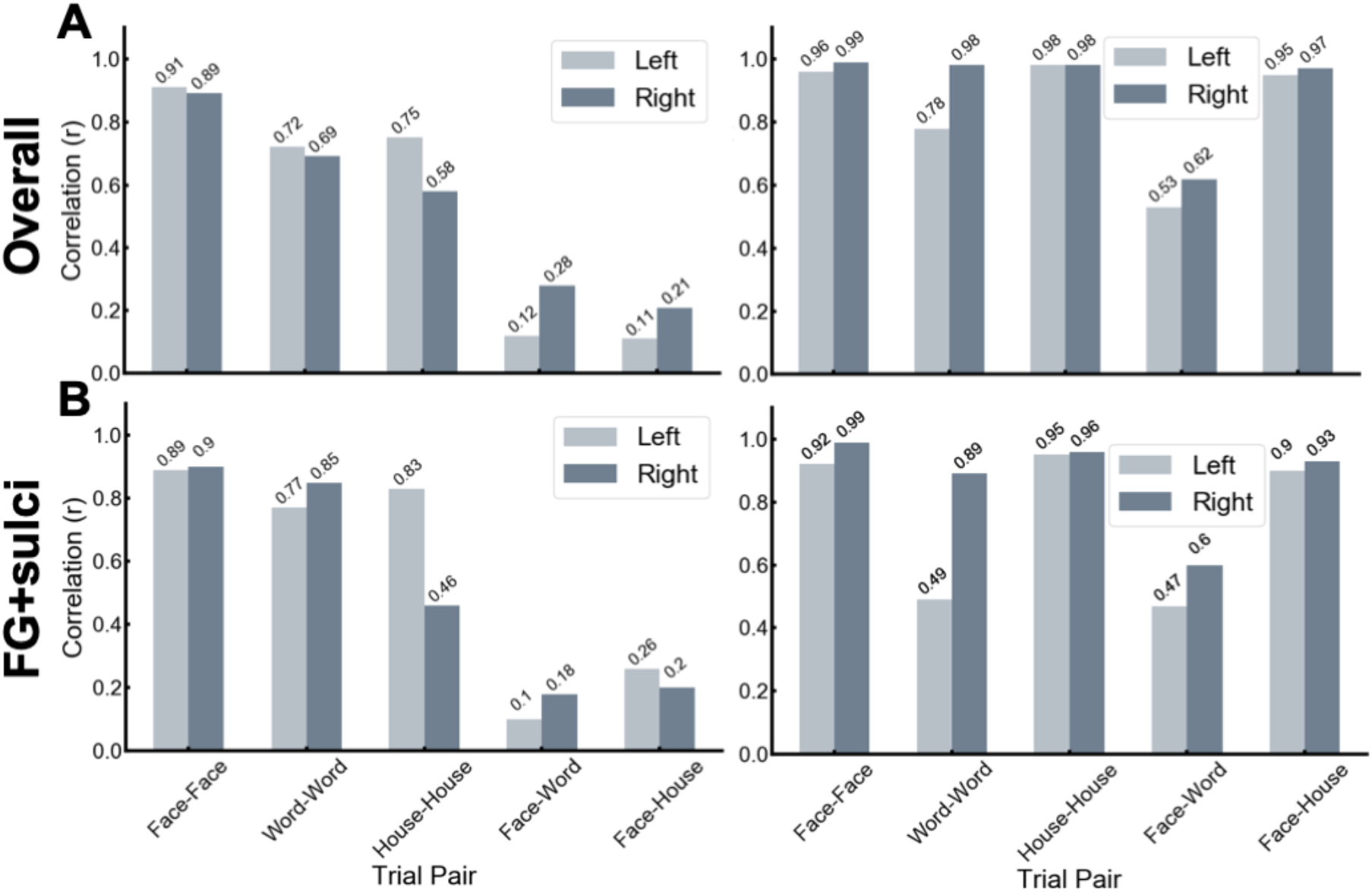
Amplitude correlations of selective responses with overlap contacts. Correlation between the face-, word- and house-selective response amplitudes within overlap contacts. **A.** Correlations across all overlap contacts, split by hemispheres, for (left) selective discrimination responses and (right) general visual responses. **B.** Correlations across all overlap contacts in the FG+sulci, split by hemispheres, for (left) selective discrimination responses and (right) general visual responses. The number on top of the bars indicates the correlation. Acronyms: FG: fusiform gyrus.

As shown in Figure 8A, in the face-word-overlap contacts, correlation between the face- and word-selective amplitudes in the left and right hemispheres was not significantly higher than 0, despite a disproportionately strong correlation for within-category amplitudes in both hemispheres. Moreover, a similar pattern was observed for faces and houses in face-house-overlap contacts (see Methods for description of correlation analysis and statistics; Left: face-word: *r* = 0.12, *p* = 0.159, 95% CI = [−0.05, 0.29]; face-face: *r* = 0.91; *p* < 0.001, 95% CI = [0.87, 0.93]; word-word: *r* = 0.72; *p* < 0.001, 95% CI = [0.63, 0.79]; face-house: *r* = 0.11; *p* = 0.235, 95% CI = [−0.07, 0.28]; house-house: *r* = 0.75, *p* < 0.001, 95% CI = [0.66, 0.82]; face-face vs. face-word: *rdiff* = 0.79; *p* < 0.001; word-word vs. face-word: *rdiff* = 0.6; *p* < 0.001; house-house vs. face-houses: *rdiff* = 0.64; *p* < 0.001. Right: face-word: *r* = 0.28; *p* = 0.096, 95% CI = [−0.02, 0.54]; face-face: *r* = 0.89; *p < 0.001*, 95% CI = [0.81, 0.94]; word-word: *r* = 0.69; *p* < 0.001, 95% CI = [0.49, 0.82]; face-house: *r* = 0.21, *p* = 0.067, 95% CI = [0.01, 0.4]; house-house: *r* = 0.58; *p* < 0.001, 95% CI = [0.43, 0.71]; face-face vs. face-word: *rdiff* = 0.61; *p* = 0.057; word-word vs. face-word: *rdiff* = 0.41; *p* = 0.012; house-house vs. face-houses: *rdiff* = 0.37; *p* = 0.039).

In contrast to the selective discrimination responses, for both the face-word-overlap and the face-house-overlap contacts, there were strong within-condition and between-condition correlations in both hemispheres, indicating that the lack of correlations in the selective discrimination responses cannot be attributed to different levels of noise or attention across conditions, since the both the general visual responses and the discrimination responses were measured concurrently within each contact (Figure 8; face-word-overlap, left: face-face: *r* = 0.96; *p* < 0.001, 95% CI = [0.95, 0.97]; word-word: *r* = 0.78; *p* < 0.001, 95% CI = [0.71, 0.84]; house-house: *r* = 0.98, *p* < 0.001, 95% CI = [0.97, 0.98]; face-word: *r* = 0.53; *p* < 0.001, 95% CI = [0.4, 0.65]; face-house: *r* = 0.95, *p* < 0.001, 95% CI = [0.93, 0.97]; Right: face-face: *r* = 0.99; *p* < 0.001, 95% CI = [0.97, 0.99]; word-word: *r* = 0.98; *p* < 0.001, 95% CI = [0.97, 0.99]; house-house: *r* = 0.98, *p* < 0.001, 95% CI = [0.97, 0.99]; face-word: *r* = 0.62; *p* < 0.001, 95% CI = [0.4, 0.78]; face-house: *r* = 0.97, *p* < 0.001, 95% CI = [0.95, 0.98]).

Finally, similar patterns were observed when considering electrode contacts only within the FG+sulci, both for the discrimination responses and the general visual responses (Figure 8B) (Discrimination responses, left: face-word: *r* = 0.1, *p* = 0.285, 95% CI = [−0.13, 0.32]; face-face: *r* = 0.89, *p* < 0.001; 95% CI = [0.82, 0.93]; word-word: *r* = 0.77, *p* = 0.001; 95% CI = [0.66, 0.85]; face-house: *r* = 0.26, *p* = 0.047, 95% CI = [0.02, 0.48]; house-house: *r* = 0.83, *p* < 0.001, 95% CI = [0.74, 0.9]; face-face vs. face-word: *rdiff* = 0.79, *p* = 0.007; word-word vs. face-word: *rdiff* = 0.67, *p* = 0.005; house-house vs. face-house: *rdiff* = 0.57; *p* < 0.001. Right: face-word: *r* = 0.18, *p* = 0.407, 95% CI = [−0.28, 0.58]; face-face: *r* = 0.90; *p* < 0.001; 95% CI = [0.77, 0.96]; word-word: *r* = 0.85, *p* < 0.001, 95% CI = [0.65, 0.94]; face-house: *r* = 0.2, *p* = 0.170, 95% CI = [−0.07, 0.43]; house-house: *r* = 0.46, *p* = 0.002, 95% CI = [0.21, 0.65]; face-face vs. face-word: *rdiff* = 0.72, *p* = 0.156; word-word vs. face-word: *rdiff* = 0.67; *p* = 0.004; house-house vs. face-house: *rdiff* = 0.26, *p* = 0.139). General visual responses: Left: face-face: *r* = 0.92, *p* < 0.001, 95% CI = [0.88, 0.95]; word-word: *r* = 0.49, *p* < 0.001, 95% CI = [0.3, 0.65]; face-word: *r* = 0.47, *p* < 0.001, 95% CI = [0.27, 0.63]; house-house: *r* = 0.95, *p* < 0.001, 95% CI = [0.92, 0.97]; face-house: *r* = 0.9, *p* < 0.001, 95% CI = [0.84, 0.94]. Right: face-face: *r* = 0.99, *p* < 0.001, 95% CI = [0.96, 0.99]; word-word: *r* = 0.89, *p* < 0.001, 95% CI = [0.74, 0.96]; face-word: *r* = 0.6, *p* = 0.011, 95% CI = [0.21, 0.82]; house-house: *r* = 0.96, *p* < 0.001, 95% CI = [0.92, 0.97]; face-house: *r* = 0.93, *p* < 0.001, 95% CI = [0.87, 0.96]).

In summary, there was no to weak correlations of between-category responses (e.g., face-word), both at the hemispheric-level and within FG+sulci, despite the same contacts showing strong within-category (e.g., face-face, word-word) correlations. Moreover, contrary to the selective responses, the general responses showed both strong within- and between-category correlations, showing that the lack of between-category correlation in the selective-responses is not due to lack of attention or disproportionate noise level. Finally, the same patterns of correlations were found for face-house-overlap contacts. Overall, these observations speak against the claim of a special relationship between faces and words in recruiting overlapping neural processes.

## Discussion

We examined the degree to which faces and written words evoke overlapping or dissociated neural responses in the human VOTC, using a large-scale intracerebral recording approach. More than two thirds of intracerebral contacts responded either *only* to faces or *only* to written words. These contacts showed strong responses in FG+sulci, with a right- and left-lateralization for faces and words, respectively, and were dissociated along the medio-lateral axis in the FG+sulci, and the posterior-anterior axis in the IOG. While the remaining 30% of contacts recorded selective responses to *both* faces and words, a similar amount of face-house-overlap was found, despite different representational demands and known neural dissociation between faces and houses. Crucially, there was little-to-no correlation in response amplitude between the face- and word-selective responses in face-word-overlap contacts, either within hemispheres or core regions of face- and word-processing (i.e., FG+sulci). Moreover, the lack of significant correlations between faces and words cannot be explained by differences in presentation frequency, attentional state, or noise, since the concurrently measured *general* visual responses showed a strong correlation, despite vastly different base stimuli. Overall, this suggest the category-*selective* human intracerebral responses to faces and written words appear largely dissociated in the human VOTC.

Studies using fMRI consistently find, especially in fusiform gyrus and surrounding sulci, substantially larger responses to faces (FFA) and words (VWFA) relative to control stimuli (Puce et al., 1995; Kanwisher et al., 1997; Cohen et al., 2000; Cohen and Dehaene, 2004; Baker et al., 2007; Davies-Thompson et al., 2016), and these functionally defined areas partially overlap in the FG+sulci (e.g., Harris et al., 2016). Similarly, ECoG studies have reported both distinct and overlapping responses to faces and written words, with a bias for overlap towards the FG (Matsuo et al., 2013). The functional relevance of this overlap is contentious as it has been proposed to reflect shared neural circuitry (e.g., Behrmann and Plaut, 2013; Robinson et al., 2017) or distinct but spatially overlapping neural circuitry (e.g., Harris et al., 2016). While fMRI studies have examined the functional relationship between the neural patterns of faces and words in VOTC regions sensitive to words (Nestor et al., 2012) or defined anatomically (Harris et al., 2016), no studies have isolated and compared the *selective* responses (i.e., faces>control vs. words>control), and there is, to our knowledge, no iEEG data to complement the (few) fMRI studies on this topic. The current study fills this gap of information by using two frequency tagging paradigms, previously used and optimized for isolating face- and word-selective responses (Rossion et al., 2015; Lochy et al., 2015), coupled with direct neural recordings from spatially sensitive intracerebral-depth-electrodes implanted in the grey matter of both sulci and gyri in a large group of patients.

The dissociations found between the selective responses to faces and written words are consistent with the claim that face and word recognition rely on spatially close yet dissociated neural circuitry. The strongest evidence supporting this claim comes from neuropsychological studies, where brain-damaged patients show highly specific impairments in the recognition of faces (prosopagnosia) or written words (pure alexia), as a result of a brain injury in the vicinity of the right and left, respectively, FG and inferior occipital gyrus (IOG) (Farah, 1991; Farah et al., 1998; Behrmann et al., 1992; Gaillard et al., 2006; Susilo et al., 2015; Robotham & Starrfelt, 2017). Moreover, neuroimaging studies consistently report enhanced face-selectivity in the right as compared to the left FG and IOG (e.g., Rossion et al., 2012; Zhen et al., 2015), while a strong left lateralization for word-selectivity is found in most studies (Cohen et al., 2000). Consistent with the view of dissociated circuitry, we found that a large proportion of the intracerebral contacts responded *only* to faces or *only* to written words, and showed the expected spatial dissociations of right and left hemispheric lateralization, respectively. Importantly, spatial dissociations were found even within core regions of face- and word-processing (FG+sulci and IOG). These findings are consistent with the recent findings of dissociated neural processes in close proximity even for the highly related tasks of reading and math (Grotheer et al., 2019).

Do overlap contacts record different processes than the above-mentioned dissociated contacts? While shared representational demands, such as need for central-view representations, could in theory lead to shared neural processes between faces and words (e.g., Hasson et al., 2002; Behrmann and Plaut, 2013; Plaut and Behrmann, 2011), we find that overlap is not unique to categories that share representational demands. That is, faces and houses showed at least a similar amount of overlap to faces and words, despite their different representational demands (central-vs. peripheral-view) and dissociated neural processes. Moreover, we found no evidence for functional relationship between face- and word-*selective* responses, despite a clear relationship in the within-category selective responses. This was in contrast to the strong between-category correlations of *general visual* responses, which were isolated at a different frequency from the *selective* responses. This latter observation may explain the comorbidity of face and word recognition difficulties following VOTC damage in terms of a general deficit (Behrmann and Plaut, 2014; Roberts et al., 2015) and the overlap of activity emphasized by some neuroimaging studies (e.g., Nestor et al., 2012). Moreover, it emphasizes the importance of isolating category-selective responses, by parsing out nonspecific visual responses, when examining functional dissociations. Thus, the present findings corroborate the view that the VOTC contain highly specialized neuro-functional circuitry for visual tasks related to face processing and reading (e.g., Harris et al., 2016; Peelen and Downing, 2017).

The existence of dissociated selective circuitry to faces and written words within the same cortical regions is still consistent with these categories being subject to competition during development for neural representation (Allison et al., 2002; Dehaene et al., 2010). FMRI studies have found evidence in favor of neural competition between faces and written words in the left VWFA, showing for example that enhancements in literacy co-occur with both an enhancement in activation to written words as well as a small decrease in activation in the same location to faces (Dehaene et al., 2010). Moreover, a 3-year longitudinal examination of the functional reorganization in a 6-year-old patient who underwent surgical removal of the right VOTC, showed a protracted increase in both face and written word selectivity in the left FG and occipito-temporal sulcus (OTS; Liu et al., 2018). Importantly, increases of face-selectivity were largely driven by the recruitment of non-selective voxels, suggesting that the competition between faces and words is largely for non-selective neuronal populations. Why do faces and written words develop in the same overall cortical regions? One possibility is that lower-level foveal retinotopic areas disproportionately project to these extrastriate areas, and that these areas are sensitive to the regularity of behaviorally relevant stimuli patterns (Levy et al., 2001; Hasson et al., 2002; Malach et al., 2002; for a discussion on cortical sensitivity to positional regularities, see Kaiser et al., 2019). An alternative explanation is that these areas contain processes crucial for encoding both face and written words, such as the encoding of larger patterns of features (Behrmann and Plaut, 2013; for recent discussion, see Op de Beeck et al., 2019). However, there are also important functional differences between faces and words. On the one hand, faces undergo little or no part decomposition in their representation (i.e., it is “holistic”, or without a category-selective representation of isolated features; Farah, 1991; Tanaka and Farah, 1993; Rossion, 2013). On the other hand, words are also represented holistically (Reicher, 1969; Farah, 1991; Pelli and Tillman, 2007) but they must also be represented in terms of their individual features (i.e., letters), which are associated to distinct sounds and typically learned independently during reading acquisition. Irrespective of the cause, the current work suggests that within the VOTC of a mature brain, face- and word-selective responses end up in close proximity but nevertheless in dissociated neuro-functional circuitry.

The present study took advantage of the high spatial resolution of depth electrodes coupled with a sensitive frequency-tagging approach that allowed for objective quantification and isolation of selective responses to faces and written words. Collectively, our findings are consistent with the view that face and word recognition in the VOTC is supported by distinct neural populations. However, given the extent of interconnectivity in the nervous system (Sherrington, 1906; Bassett and Sporns, 2017), research should examine the connections between different specialized circuitry, as they could potentially modulate (e.g., inhibit) each other. Indeed, ECoG recordings, although limited in its amount of data (4 patients), suggest a mechanism for face-selective circuitry to modulate word-selective circuitry, but not vice versa (Matsuo et al., 2013; see also Allison et al., 2002). Finally, an open question that deserves further attention is whether dissociated circuitry is specific to highly specialized skills acquired relatively early in life – such as face detection, reading, and spatial navigation – or whether specialization in any visual domain even in later stages of life ultimately leads to specialized and dissociated circuitry. Indeed, dissociation of neural circuitry could be a fundamental neural organizing principle of the brain for organizing quick and accurate stimulus-response mappings.

## Acknowledgements

The research has been funded by a LUE (Lorraine Université d’Excellence) program, a 2018 project from the Région Grand Est, and a FNRS EOS project (ID: 30991544).

